# CALFP-MHC: Interpretable Pan-Allelic Prediction of Peptide-MHC Binding and Presentation Using Chemically Grounded Fingerprints and Contrastive Learning

**DOI:** 10.64898/2026.08.10.743877

**Authors:** My-Diem Nguyen Pham, Thi-Kim-Cuong Ho, Hoai-Nghia Nguyen, Le-Son Tran, Minh-Duy Phan, Vy Nguyen

## Abstract

Identifying which peptides bind major histocompatibility complex (MHC) molecules is central to vaccine design, neoantigen prioritization, and precision immunotherapy. Existing deep learning predictors largely encode amino acids as discrete symbols, thereby missing the residue-level chemistry driving molecular recognition. Performance also tends to degrade under class imbalance, for rare alleles, and on peptide– MHC combinations outside the training distribution.

We developed CALFP-MHC, a framework that encodes each amino acid as a set of complementary cheminformatics fingerprints capturing functional groups, atomic connectivity, and substructural features, and combines positional encoding with supervised contrastive pre-training to organize the latent space by binding class before fine-tuning a binary classifier. Peptide–MHC interactions are modeled through a hybrid convolutional-transformer backbone.

In a large-scale computational benchmark covering ∼18.7 million peptide–MHC pairs across 112 HLA class I and 53 class II alleles, CALFP-MHC achieved AUCs of 0.93-0.97 and PPVs of 0.66–0.94. Critically, performance remained above AUC 0.90 even at a 200:1 negative-to-positive ratio, where competing tools frequently collapsed toward chance. On independent experimental data containing 3,627 class I and 520 class II MS/MS-confirmed ligands and 570 validated neoantigens, the model maintained strong discrimination, correctly prioritizing immunogenic peptides and MHC-presented ligands. Attention and integrated-gradient analyses recovered established anchor positions (P2 and PΩ for class I, P1, P4, P6, and P9 for class II) and highlighted chemically interpretable functional groups consistent with known binding determinants.

CALFP-MHC demonstrates that grounding residue representations in molecular chemistry, rather than sequence symbols alone, improves both robustness and interpretability in peptide–MHC binding prediction.

## Introduction

Prediction of peptide–MHC (pMHC) binding is a fundamental task in computational immunology, with applications in neoantigen discovery, vaccine design, and autoimmune disease research. Accurate identification of binding peptides from large candidate pools is essential because only a small fraction of mutated peptides is ultimately presented to T cells and recognized as immunogenic.

Although several high-performing prediction models, including NetMHCpan^1^, MHCflurry^2^, BigMHC^3^, MixMHCpred^4^, and DeepMHCII^5^, have achieved strong predictive performance, important challenges remain. Existing methods represent amino acids as discrete symbols^1,5,6^ or one-hot vectors^2,3,7^, which inadequately capture the physicochemical properties of residue side chains that govern molecular recognition and peptide–MHC binding. In addition, model performance often declines substantially under conditions of severe class imbalance, for rare alleles, and for peptide–MHC pairs that fall outside the training distribution. Many current approaches^7,8^ also provide limited interpretability, offering little insight into the specific residues or chemical features responsible for binding decisions. This lack of explainability restricts their utility in mechanistic studies and translational applications, where biological interpretation is as important as predictive accuracy.

Several directions have been explored to address these limitations. Regarding residue representation, Lee and Min (2023)^9^ proposed repurposing molecular fingerprints for amino acid encoding, and Adamczyk *et al*. (2026)^10^ demonstrated that MACCS keys (Molecular ACCess System, 167-bit), ECFP4 (Extended-Connectivity Fingerprints at radius 2, 1,024-bit), ECFP6 (at radius 3, 2,048-bit), and RDKit fingerprints (1,024-bit) at the amino acid level achieve competitive performance across peptide benchmarks, suggesting that cheminformatics-based encodings can capture side-chain chemistry more explicitly than discrete or one-hot representations. In parallel, contrastive learning has shown promise for organizing binding-relevant latent representations: ConBoTNet^11^ applied supervised contrastive pre-training with a bottleneck transformer specifically for MHC-II peptide binding. Nevertheless, no existing approach integrates chemically informed residue representations, explicit peptide–MHC interaction modeling, and joint prediction of both binding affinity and antigen presentation across MHC class I and class II systems.

We hypothesized that representing amino acids using molecular fingerprints encoding functional groups, atomic connectivity, and substructural chemical features would provide a more faithful description of the biochemical determinants underlying peptide–MHC binding. To test this hypothesis, we developed CALFP-MHC, a deep learning framework that represents each amino acid using four complementary molecular fingerprint descriptors (MACCS keys, ECFP4, ECFP6, and RDKit), integrates positional information through trigonometric encoding^12^, and models peptide–MHC interactions using a hybrid convolutional–transformer architecture. The framework employs a two-stage training strategy in which supervised contrastive learning is first used to organize the latent representation space according to binding class, followed by fine-tuning of a binary classifier for affinity prediction. We evaluated CALFP-MHC across 112 HLA class I alleles (48 used for head-to-head benchmarking) and 53 class II alleles under challenging conditions, including severe class imbalance, rare alleles, and previously unseen peptides. Across these benchmarks, CALFP-MHC consistently outperformed existing prediction methods in terms of AUC and positive predictive value while also generating interpretable binding motifs that aligned with known structural interaction patterns.

## Results

### Overview of CALFP-MHC

CALFP-MHC was designed to capture key topological and structural features of peptide–MHC interactions while preserving residue-level positional information. Peptides and MHC pseudo-sequences (34-residue contact-position representations as defined by Nielsen et al^13,14^) were encoded using four descriptor types: MACCS, ECFP4, ECFP6 and RDKit fingerprints. Each MHC pseudo-sequence comprises the 34 polymorphic amino acid positions located within 4.0 Å of the bound peptide in representative MHC crystal structures (see Methods). MACCS keys (167-bit) encode predefined functional groups (e.g., nitrogen-containing, hydroxyl and aromatic moieties) relevant to molecular recognition, including hydrogen bonding and hydrophobic interactions. ECFP4 (1,024-bit) and ECFP6 (2,048-bit) fingerprints represent local atomic environments and intra-residue connectivity at radii of 2 and 3 bonds, respectively. RDKit fingerprints (1,024-bit) capture linear bond connectivity within residues. Collectively, these descriptors provide a compact representation of functional groups, substructures and topological connectivity in peptides and MHC molecules (Fig. 1b).

**Fig. 1.**
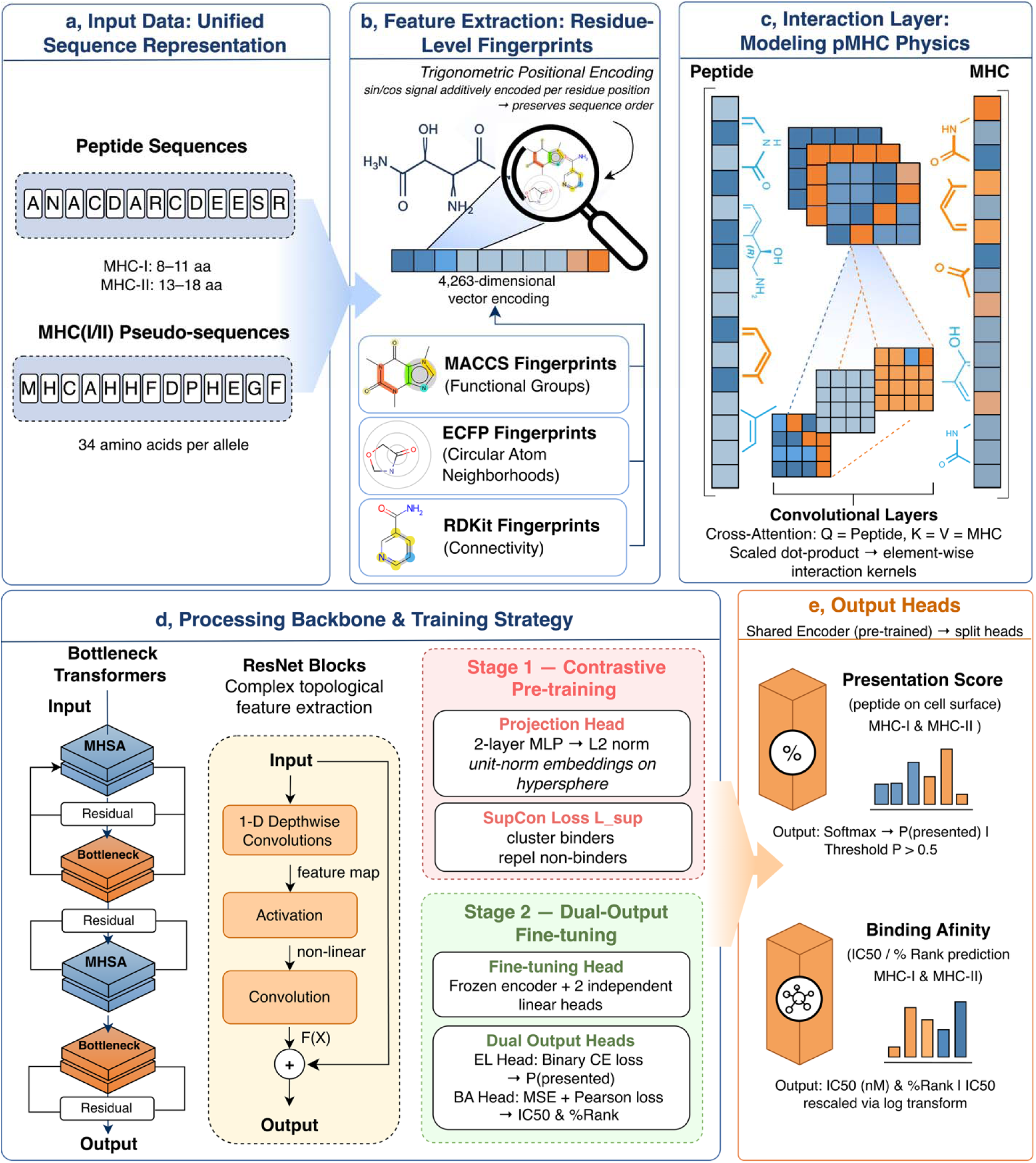
| Fingerprint-based deep learning framework for peptide–MHC binding prediction with dual output heads. **a**, Peptide sequences (MHC-I: 8–11 aa; MHC-II: 13–18 aa) and MHC(I/II) pseudo-sequences (34-residue representations derived from polymorphic positions lining the peptide-binding groove, as defined by Nielsen et al ^13,14^) are provided as input. **b,** Each residue is encoded as a 4,263-dimensional fingerprint vector by concatenating MACCS (166-bit), ECFP (1,024-bit), ECFP6 (2,048-bit), and RDKit (1,024-bit) fingerprints; sinusoidal positional signals are additively encoded per residue position to preserve sequence order. **c,** Peptide and MHC fingerprint matrices interact via cross-attention (Q = Peptide, K = V = MHC) followed by convolutional layers to model element-wise physical and chemical interactions. **d,** Interaction features are processed by a bottleneck transformer (multi-head self-attention with residual connections) combined with ResNet blocks (1-D depth-wise convolutions with identity skip connections) for high-level feature refinement. Training proceeds in two stages: Stage 1 applies supervised contrastive learning (SupCon loss) with a 2-layer MLP projection head to cluster binders and repel non-binders in a unit-norm embedding space; Stage 2 fine-tunes a frozen encoder with two independent linear heads — an EL head (binary cross-entropy loss → presentation probability) and a BA head (MSE + Pearson correlation loss → IC50 and %Rank). **e,** Output heads produce a Presentation Score (softmax → P(presented), threshold P > 0.5) and Binding Affinity (IC50 in nM and %Rank, IC50 rescaled via log transform) for both MHC-I and MHC-II.

To incorporate residue positional information, each fingerprint vector was combined with a sinusoidal positional encoding via element-wise addition (see Methods), where *p* denotes the position of the residue in the peptide or MHC sequence. This transformation assigns position-dependent weights to fingerprint features, enabling identical amino acids appearing at different sequence locations to contribute differentially to the interaction. By integrating compositional and positional information, this approach provides a more expressive representation of peptide–MHC interactions.

Using these input features, we developed a hybrid architecture that combines convolutional layers to capture local peptide–MHC interactions with transformer attention layers to model long-range dependencies along the binding interface (Fig. 1c–d). To optimize performance, we adopted a two-stage training strategy. First, supervised contrastive learning structures the latent space by clustering binding pairs while separating non-binding pairs, improving generalization to unseen peptides and MHC alleles (Fig. 1d – Stage 1). In the second stage, the model is fine-tuned with a binary classifier to enhance predictive accuracy (see Methods). Beyond prediction, CALFP-MHC provides interpretability through attention weights that highlight residues contributing to the binding core (Fig. 1d – Stage 2).

Using the same framework, we developed CALFP-MHCI and CALFP-MHCII to model peptide interactions with MHC class I and MHC class II, respectively. The models differ primarily in peptide input length, reflecting the shorter peptides presented by MHCI (8–11 amino acids) compared with those bound by MHCII (typically 13–18 amino acids). CALFP-MHCI and CALFP-MHCII are therefore evaluated separately.

### CALFP-MHCI outperformed existing tools in predicting the binding between HLAI peptide

CALFP-MHCI was benchmarked against five state-of-the-art pMHC binding predictors (NetMHCpan, BigMHC, deepAntigen, MixMHCpred and MHCflurry) across 48 HLA class I alleles (Supplementary Table S1), comprising 14 HLA-A, 21 HLA-B and 13 HLA-C. The number of peptide–HLA pairs varied widely across alleles, ranging from 45 to 439,272. Overall, CALFP-MHCI showed consistently strong performance, with AUC values typically between ∼0.95 and 1.0 and PPV generally above 0.8 (Fig. 2a). In comparison, other methods exhibited greater variability, with some alleles (HLA-B*39:10, HLA-B*35:04 and HLA-B*44:05) approaching near-random performance (AUC ∼0.5–0.6; PPV < 0.3). Together, these results suggest that CALFP-MHCI is robust across alleles with diverse data availability (Fig. 2a).

**Fig. 2.**
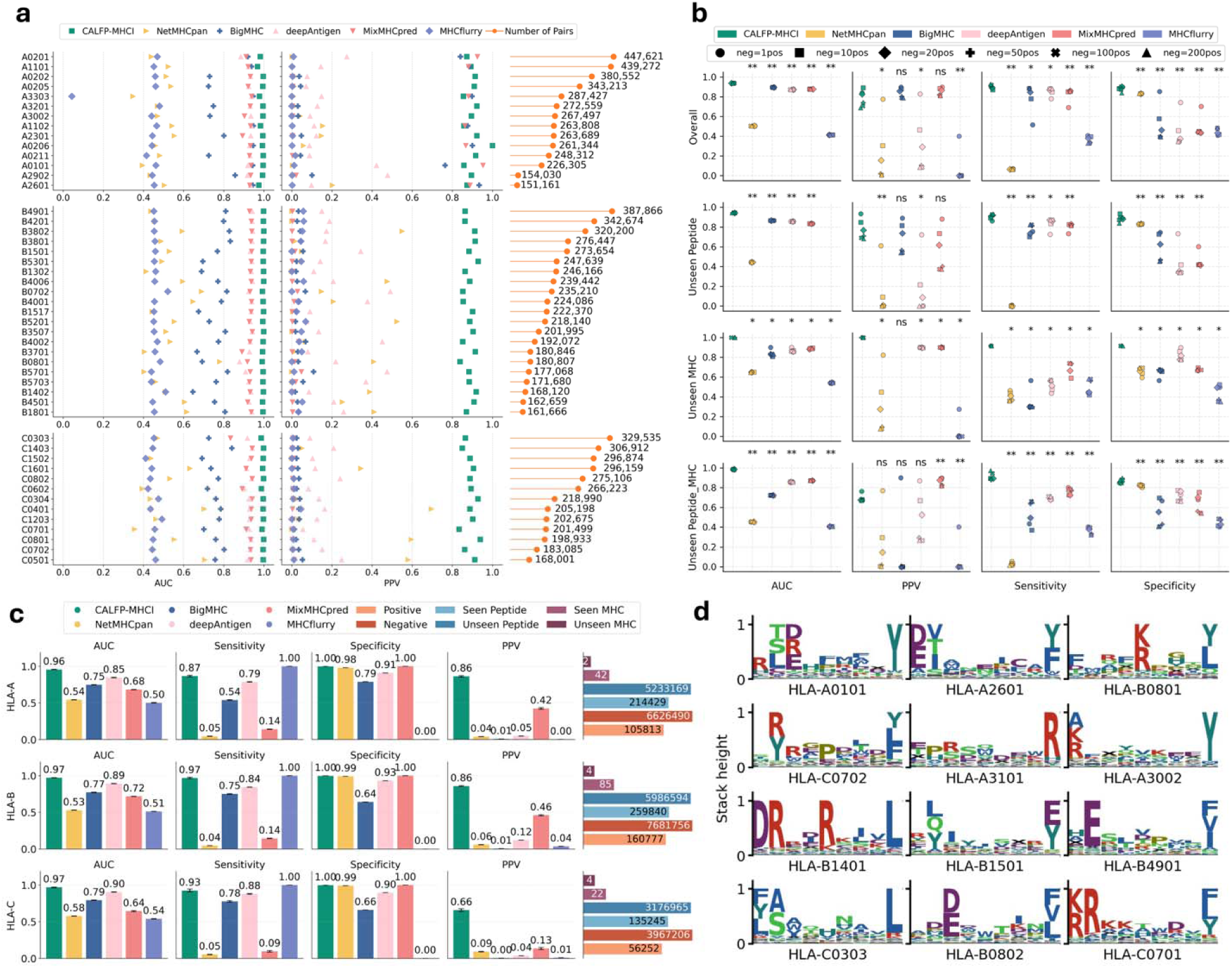
| Benchmarking and interpretability of the fingerprint-based peptide–MHC class I model. **a**, Allele-specific benchmarking showing consistent performance of CALFP-MHCI across representative HLA class I alleles. **b,** Comparison with state-of-the-art predictors under varying negative sampling ratios, evaluated by AUC, PPV, sensitivity, and specificity. **c,** Per-allele metrics highlighting improved performance of CALFP-MHCI relative to existing methods. Counts of positive, negative, seen peptide, unseen peptide, seen MHC, and unseen MHC in the testing dataset are shown in the right bar plot. **d,** The y-axis represents relative residue enrichment at each position, with stack height reflecting the degree of amino acid conservation; taller stacks indicate stronger positional preference.

On the same dataset, we also assessed CALFP-MHCI across HLA-A, HLA-B, and HLA-C loci (Fig. 2c). CALFP-MHCI maintained consistently high performance across all loci, with mean AUCs of 0.96 (HLA-A), 0.97 (HLA-B) and 0.97 (HLA-C), compared with 0.50–0.79 for other methods. The sensitivity and specificity remained high (0.93–1.0), whereas competing models often showed reduced sensitivity (<0.2) or specificity (<0.7). The positive predictive values were 0.86 for HLA-A, 0.86 for HLA-B, and 0.66 for HLA-C, suggesting that CALFP-MHCI is robust to unseen peptides and shows reasonable generalization to less-represented MHC molecules.

We further evaluated performance across datasets with negative:positive sampling ratios ranging from 1:1 to 200:1. CALFP-MHCI remained relatively stable, maintaining AUC values above 0.9 and PPV of ∼0.8–0.9. It should be noted that CALFP-MHC was trained on a balanced dataset (1:1 ratio), whereas competing tools were trained on their own data distributions; the observed robustness under imbalance may partly reflect this training difference. Nevertheless, other predictors showed marked performance declines; at a 100:1 ratio, NetMHCpan and BigMHC fell below 0.6, whereas deepAntigen, MixMHCpred and MHCflurry approached near-random performance (PPV ≤ 0.2) (Fig. 2b). This trend was also observed for unseen peptides and MHC molecules, where CALFP-MHCI exceeded competing methods by ∼0.3–0.4 in both AUC and PPV.

Beyond predictive performance, CALFP-MHCI yielded interpretable binding motifs through attention-based analysis (Fig. 2d). The resulting sequence logos recovered established anchor preferences, such as tyrosine (Y) at the C terminus for HLA-A*01:01 and HLA-A*26:01, and arginine (R) at position 5 for HLA-B*08:01^15,16^. For well-characterized alleles (including HLA-A*01:01, HLA-A*26:01 and HLA-B*08:01), the inferred motifs were broadly consistent with those reported by MixMHCpred and PRIME. Notably, for less-characterized alleles (for example, HLA-C*07:01, HLA-C*03:03 and HLA-B*08:02), CALFP-MHCI suggested additional motif features, such as enrichment of arginine (R) at positions P1 and P2 in HLA-C*07:01, which are less consistently captured by other predictors. These results indicate that CALFP-MHCI can recover known binding preferences while providing potentially informative insights into peptide–MHC recognition.

### CALFP-MHCII outperformed existing tools in predicting the binding between HLAII-peptide

We next evaluated CALFP on MHC class II alleles, which present peptides to CD4 T cells and exhibit distinct binding properties owing to their open-ended binding groove and broader peptide-length diversity.

CALFP-MHCII was benchmarked against five state-of-the-art predictors (NetMHCIIpan^17^, DeepMHCII^5^, DeepSeqPanII^7^, MixMHC2pred^6^ and TLimmuno2^18^) across a comprehensive panel of HLA class II alleles (Supplementary Table S2). At the allele level (Fig. 3a), CALFP-MHCII showed consistentl strong performance, with AUC values typically approaching 0.9–1.0 and PPVs generally above 0.7, whereas competing methods exhibited greater variability (AUC 0.50–0.88; PPV 0.00–0.69). Th performance gap was more pronounced for alleles with limited training data, such as HLA-DPB1*15:01 (n = 34 pairs), where most competing methods approached random performance (AUC = 0.50, PPV = 0.00 for DeepSeqPanII, MixMHC2pred, NetMHCIIpan, and TLimmuno2), whilst CALFP-MHCII achieved AUC = 0.87 and PPV = 0.94. For well-represented alleles, including HLA-DRB1*04:01, CALFP-MHCII maintained high performance (AUC = 0.95, PPV = 0.93), suggesting robustness across both data-sparse and data-rich settings.

**Fig. 3.**
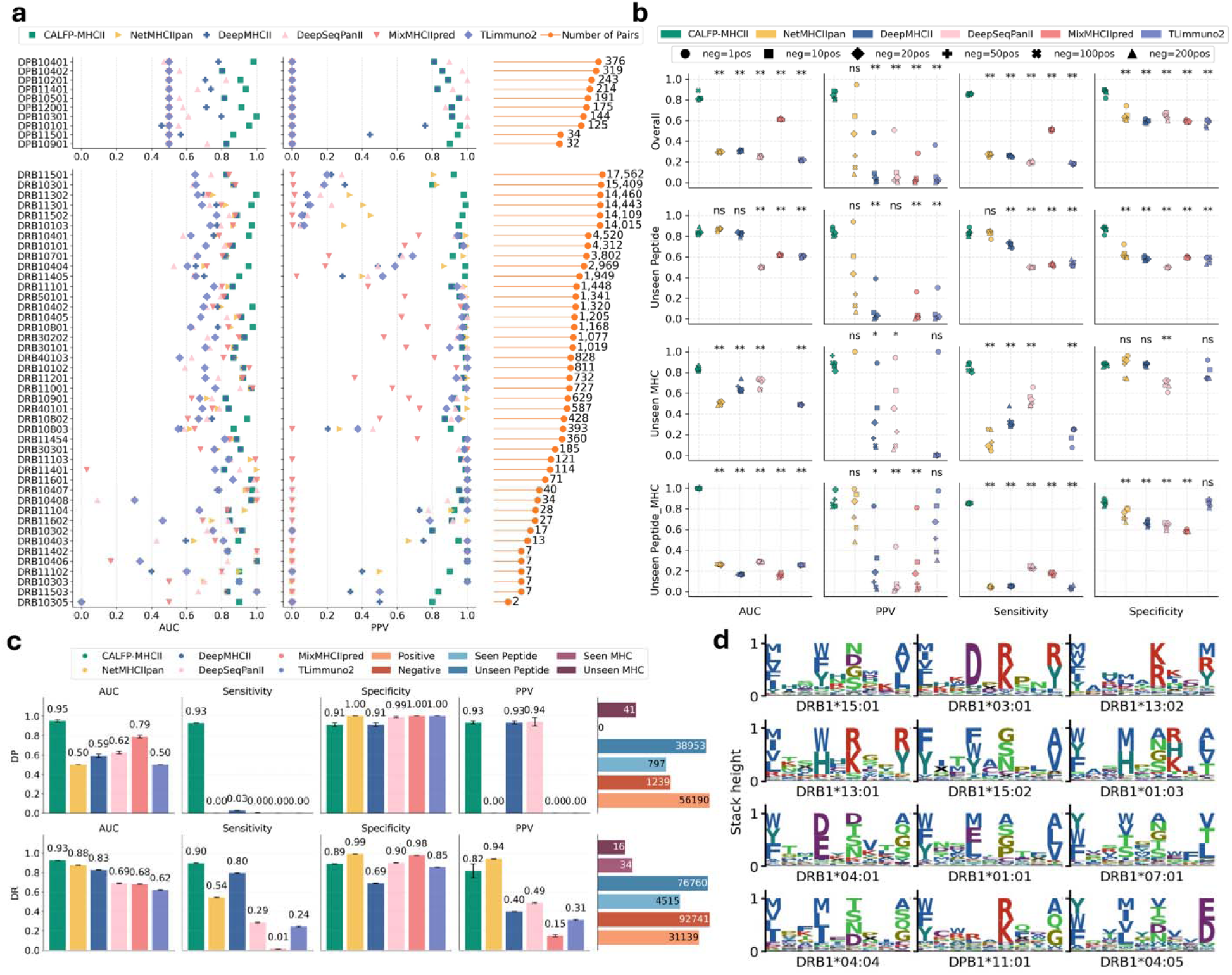
| Performance and interpretability of CALFP-MHCII for peptide–MHC class II binding prediction. **a**, Allele-specific benchmarking across diverse HLA class II molecules, showing consistent improvements of CALFP-MHCII i AUC and PPV over existing predictors. **b,** Comparison with state-of-the-art methods under varying negative sampling ratios, evaluated by AUC, PPV, sensitivity, and specificity. **c,** Per-allele benchmarking highlighting superior performance of CALFP-MHCII across representative alleles. **d,** Attention-derived sequence logos illustrating allele-specific binding motifs captured by the model.

In the negative:positive imbalance benchmark (Fig. 3b), CALFP-MHCII, being trained on a balanced dataset (1:1 ratio), maintained stable performance, with AUC values above 0.9 and PPV of ∼0.7–0.8 as the ratio increased from 1:1 to 200:1. In contrast, other models showed marked performance declines, with PPVs often below 0.3 and AUCs approaching near-random levels (∼0.5) at higher imbalance ratios (from 50:1 to 100:1).

On the same dataset, we assessed model generalization across three challenging scenarios: interaction of unseen peptides, or unseen MHC, and their combination (Table S1). Across these scenarios, CALFP-MHCII consistently outperformed the strongest baseline by ∼0.2–0.4 in both AUC and PPV, indicating improved generalization. At the locus level (Fig. 3c), CALFP-MHCII maintained robust performance for HLA-DP (AUC 0.95, sensitivity 0.93, specificity 0.91, PPV 0.93) and HLA-DR (AUC 0.93, sensitivity 0.90, specificity 0.89, PPV 0.82), whereas competing methods often showed reduced sensitivity (<0.2) or lower PPV (∼0.3–0.5).

We assessed interpretability of CALFP-MHCII by extracting attention-based sequence logos (Fig. 3d). CALFP-MHCII recovered motifs consistent with established biochemical preferences, including enrichment of arginine (R) at position P6 for HLA-DRB1*03:01 and HLA-DRB1*13:02, hydrophobic residues at position P1 for HLA-DRB1*01:03 (methionine, M) and HLA-DRB1*07:01 (tyrosine, Y), enrichment of basic residues (arginine, R and lysine, K) at position P6 for HLA-DPB1*11:01. CALFP-MHCII also suggested informative motifs for less-represented alleles, such as HLA-DPB1*04:01 and HLA-DRB1*04:05, including enrichment of aspartic acid (D) at position P4 and glutamic acid (E) at the C-terminal anchor position, respectively. Compared with MixMHC2pred and PRIME, CALFP-MHCII recapitulated known motifs for well-studied alleles while identifying additional sequence features, supporting both its predictive performance and interpretability.

### Fingerprint-based contrastive learning enables interpretable prediction of MHC–peptide interactions

To evaluate the interpretability of CALFP-MHC across multiple levels, we analyzed positional amino acid preferences, structural contacts, chemical fingerprints, internal attention, and latent space organization (Fig. 4a–k). At the sequence level, amino acid preference maps (Fig. 4a, c) revealed that the model learned canonical anchor motifs for both class I and class II peptides. For class I (Fig. 4a), the model recovered allele-specific anchor patterns consistent with known binding preferences^15,16^: P2 for HLA-B*44:02 and PΩ for HLA-A*68:01. For MHC class II (Fig. 4c), the model also recovered allele-specific patterns: P4 was the dominant anchor position for HLA-DPA1*02:01/DPB1*01:01 and P8 for HLA-DRB1*03:01, consistent with widely recognized anchor positions within the MHC-II binding groove^6,21^. These anchor residues map directly onto the MHC binding pockets in the corresponding crystal structures (Fig. 4b, d): P2 and PΩ occupy the hydrophobic B and F pockets in class I, while the identified class II positions form hydrogen bonds and electrostatic interactions within the binding groove^19,21^, confirming that the motifs captured by CALFP-MHC are consistent with physically grounded peptide–MHC interactions.

**Fig. 4.**
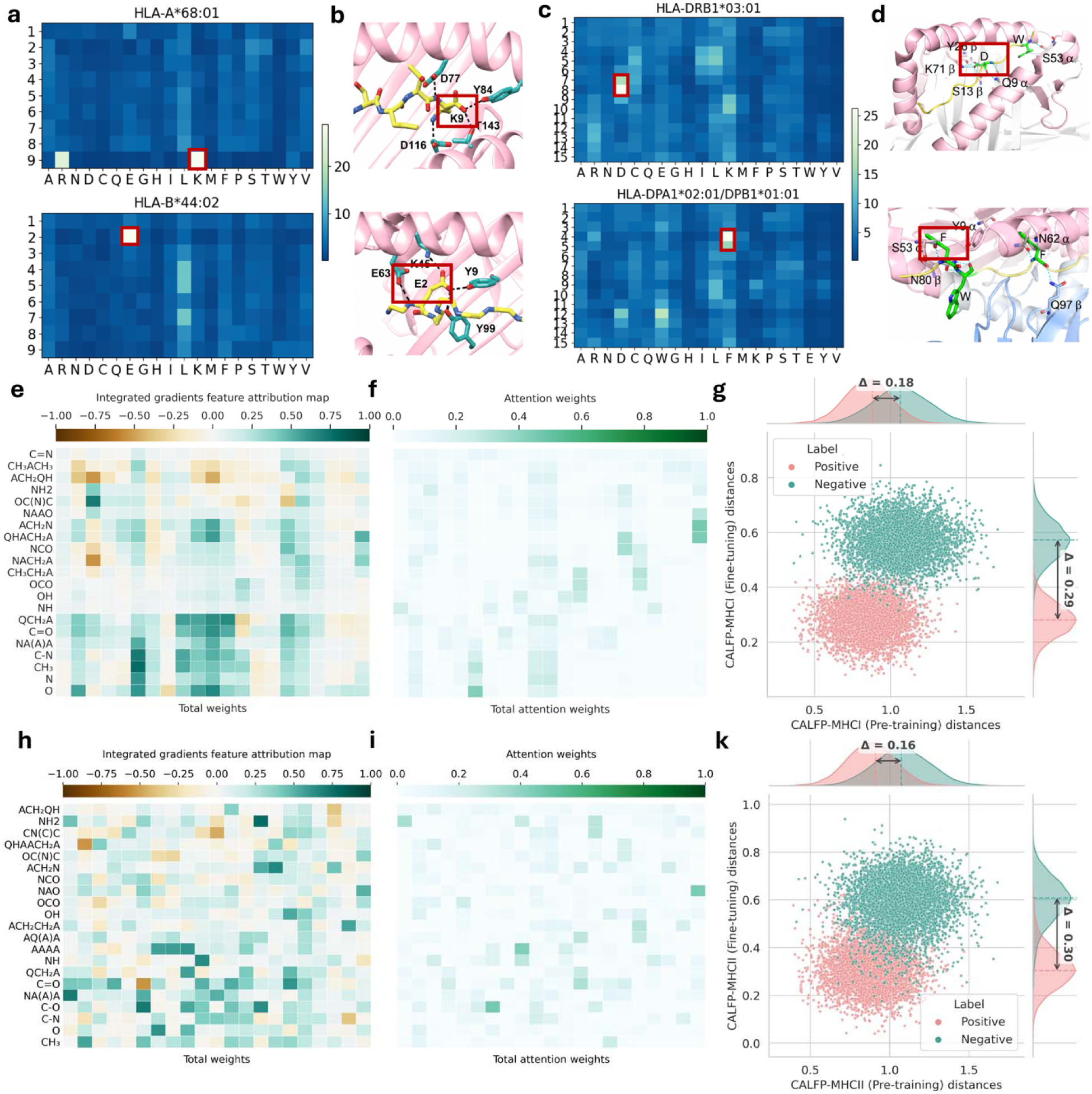
| Fingerprint-based contrastive learning enables interpretable prediction of peptide–MHC interactions. **a, c**, Heatmaps of positional amino acid preferences showing residue-level anchor patterns for class I. Each column corresponds to a peptide position and each row to one of the 20 amino acids; brighter pixels indicate higher accumulated attention scores. **b, d,** Structural data supporting the anchor motifs in panels a and c, respectively. Red boxes mark the dominant anchor residue in each heatmap **(a, c)** and the corresponding residue in the matched structural view **(b, d)**: PΩ (K9) for HLA-A*68:01 and P2 (E2) for HLA-B*44:02, mapped onto a 9-mer peptide–HLA-A*68:01 complex (PDB: 4HWZ); and P8 (D) for HLA-DRB1*03:01 and P4 (F) for HLA-DPA1*02:01/DPB1*01:01, mapped as a representative example onto CII bound to HLA-DRB1*01:01 (PDB: 2FSE)^6,19,20^. **e, h,** Integrated gradients attribution maps highlighting fingerprint substructures contributing to binding in class I **(e)** and class II **(h)**. **f, i,** Attention weight heatmaps indicating residue-level focus on informative chemical features. **g, k,** Latent-space distance distributions for MHCI **(g)** and MHCII **(k)** after supervised contrastive fine-tuning, showing increased inter-class separation (MHCI: Δ = 0.29; MHCII: Δ = 0.30) and global restructuring of the embedding space.

At a deeper level, integrated gradients attribution maps (Fig. 4e, h) highlighted functional groups such as amines, carbonyls, hydroxyls, and aromatic rings, which are known to mediate hydrogen bonding, charge stabilization, and hydrophobic packing^20^. Attention weight maps (Fig. 4f, i), which should be interpreted with caution as attention weights are not guaranteed to reflect feature importance, nonetheless showed broadly overlapping hotspots at the same anchor positions identified by integrated gradients attribution, particularly at P2 and PΩ for class I / P4 and P6 for class II, with elevated weights concentrated in (ECFP-based / MACCS) fingerprint dimensions encoding (aromatic and charged) substructures. The co-localization of attribution and attention signals at these positions indicates that CALFP-MHC internally prioritizes chemically relevant features rather than distributing weight uniformly across the sequence. This consistency between model-internal attention and post hoc attribution suggests that CALFP-MHC focuses on meaningful chemical determinants, thereby strengthening its reliability and interpretability.

We further demonstrate the strong performance of CALFP-MHC at the representation level. Latent distance distributions before and after fine-tuning (Fig. 4g, k) show a pronounced reorganization of the embedding space. Before fine-tuning, positive and negative pairs largely overlapped, limiting separability. After supervised contrastive learning, positive pairs contract to smaller distances, while negative pairs shift outward, reducing overlap across the distribution. This change indicates a global restructuring of the embedding manifold rather than local refinements.

Taken together, the consistency across sequence motifs (Fig. 4a, c), structural mapping (Fig. 4b, d), chemical attribution (Fig. 4e, h), attention weights (Fig. 4f, i), and latent embeddings (Fig. 4g, k) demonstrates that CALFP-MHC is consistent with prioritizing chemically and structurally relevant features, though direct experimental validation of these attributions remains an important future direction. Therefore, CALFP-MHC demonstrated strong predictive performance and provided interpretable mechanistic insight into peptide–MHC recognition, supporting its use in immunology.

### CALFP-MHC accurately identifies MHC-presenting peptides across experimental datasets

We next evaluated CALFP-MHC on independent experimental datasets to assess its ability to identify MHC-presented peptides across distinct assay modalities (Fig. 5a–c). For MHCI, an MS/MS dataset of HLA-presented ligands was collected from a previous study (Fig. 5a), containing 2,827 positives and 800 negatives, with pronounced allele imbalance: a subset of alleles is well represented, whereas others remain sparsely sampled^22^. For class II, we assembled a composite dataset integrating MS/MS-identified ligands and IC –based binding assays (Fig. 5b)^17^, comprising 306 positives and 214 negatives, with most peptides restricted to DRB1, DQA, and DPB alleles. Together, these complementary cohorts span distinct assay modalities and allele distributions, providing a stringent framework for external validation.

**Fig. 5.**
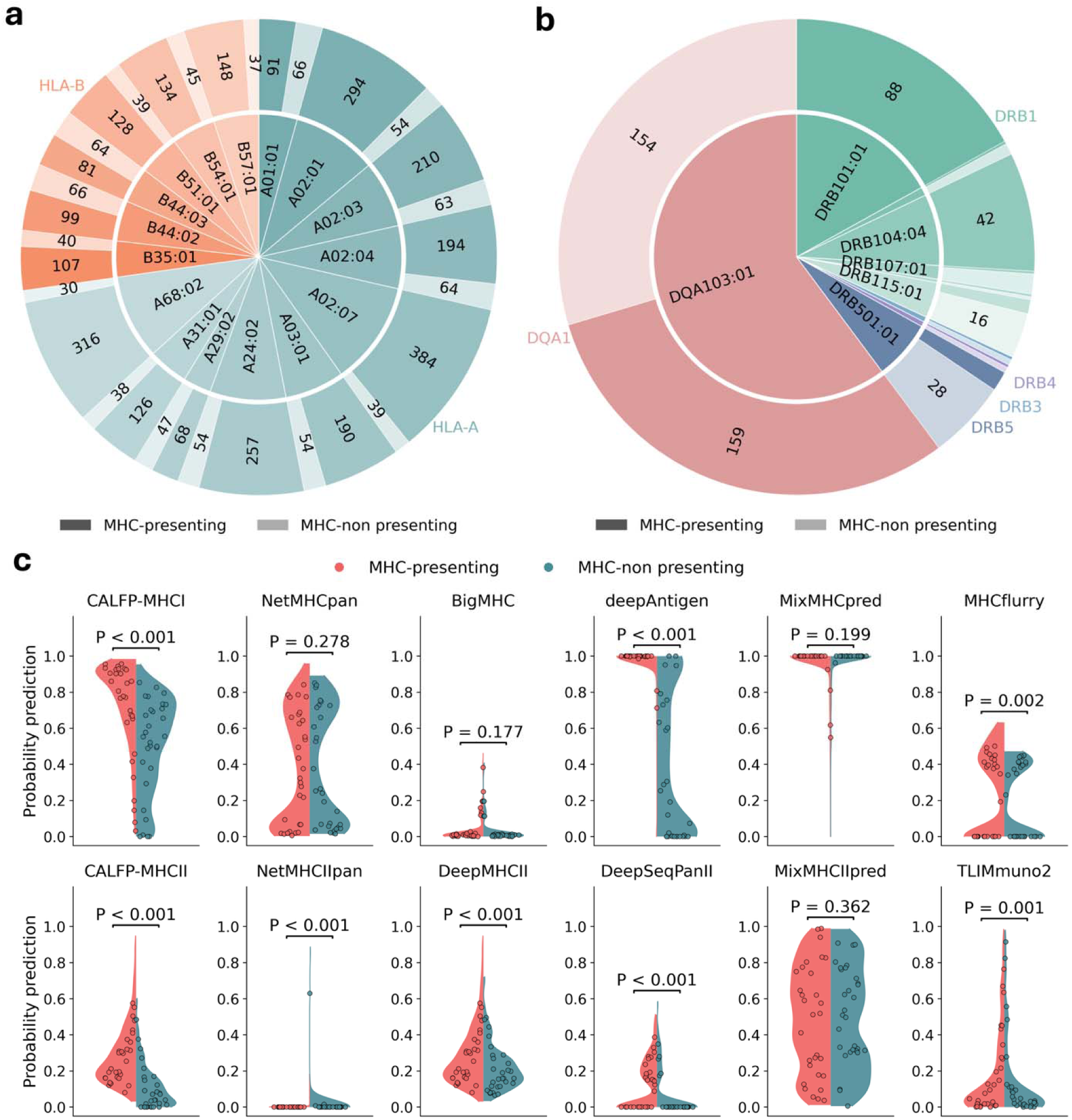
| CALFP-MHCI and CALFP-MHCII identify MHC-presenting peptides from independent experimental datasets. **a**, Allele-specific distribution of MHC-presenting and non-presenting peptides across HLA class I alleles in the MS/MS-validated dataset²². Each donut chart shows the overall proportion of MHC-presenting (inner ring) and non-presenting peptides, with allele-specific counts in the outer ring. **b,** Allele-specific distribution of MHC-presenting and non-presenting peptides across HLA class II alleles from a composite dataset integrating MS/MS ligands and IC binding assays^17^. **c,** Prediction probability distributions for MHC-presenting (positive) and MHC-non-presenting (negative) peptides. Top row shows predictions on the class I MS/MS dataset **(a)**; bottom row shows predictions on the class II dataset **(b)**. Six predictors are shown per row in the same layout and colour scheme. Statistical comparisons were performed using the Mann–Whitney U test (two-sided); p-values are shown above each violin.

Across these datasets, CALFP consistently achieved better separation between positive and negative peptides compared with state-of-the-art predictors. In the class I MS/MS dataset (Fig. 5c), CALFP-MHCI assigned higher probabilities to validated ligands and compressed negatives toward lower scores, yielding a strikingly reduced overlap and highly significant separation (p < 0.001). By contrast, tools such as NetMHCpan^1^, MixMHCpred^4^, and MHCflurry^2^ produced broader overlaps. In the class II composite cohort (Fig. 5c), CALFP-MHCII similarly achieved clear discrimination (p < 0.01), outperforming DeepSeqPanII^7^, DeepMHCII^5^, and TLImmuno2^18^, with distributions of positives and negatives distinctly separated.

### CALFP-MHC prioritizes immunogenic neoantigens for personalized cancer treatment

Having established strong performance in identifying MHC-presented peptides, we next asked whether CALFP-MHC could further distinguish immunogenic from non-immunogenic peptides, a more stringent challenge relevant to personalized cancer treatment. To this end, we used the TESLA (Tumor Epitope Selection by Lymphocyte Assessment) dataset^23^, an independent collection of candidate neoantigens with experimentally validated T cell recognition. The non-immunogenic neoantigens substantially outnumber the immunogenic ones (36 immunogenic versus 534 non-immunogenic after preprocessing), reflecting the scarcity of validated neoantigens.

CALFP-MHCI achieved the strongest discrimination between immunogenic and non-immunogenic neoantigens (p < 0.001), whereas NetMHCpan showed no significant separation (p = 0.980), and BigMHC remained non-significant (p = 0.086). We note that the TESLA dataset^23^ contains only 36 immunogenic peptides after preprocessing, which limits statistical power; the dataset is adequately powered to detect large discrimination effects (AUC ≥ 0.65) but cannot reliably rank predictors whose true AUC difference is smaller than approximately 0.10-0.15. The reported p-values reflect discrimination ability rather than superiority over competing methods. Although deepAntigen and MHCflurry^2^ also reached significance (p < 0.001), their score distributions showed broader overlap between positive and negative peptides compared with CALFP-MHCI. Although failing to discriminate on the MS/MS dataset (p = 0.199, Fig. 5c), MixMHCpred achieved significance on the validated neoantigen dataset (Fig. 6b), suggesting inconsistent performance across evidence types. Together, these results suggest that CALFP-MHC captures predictive features of T cell recognition beyond MHC presentation alone.

**Fig. 6.**
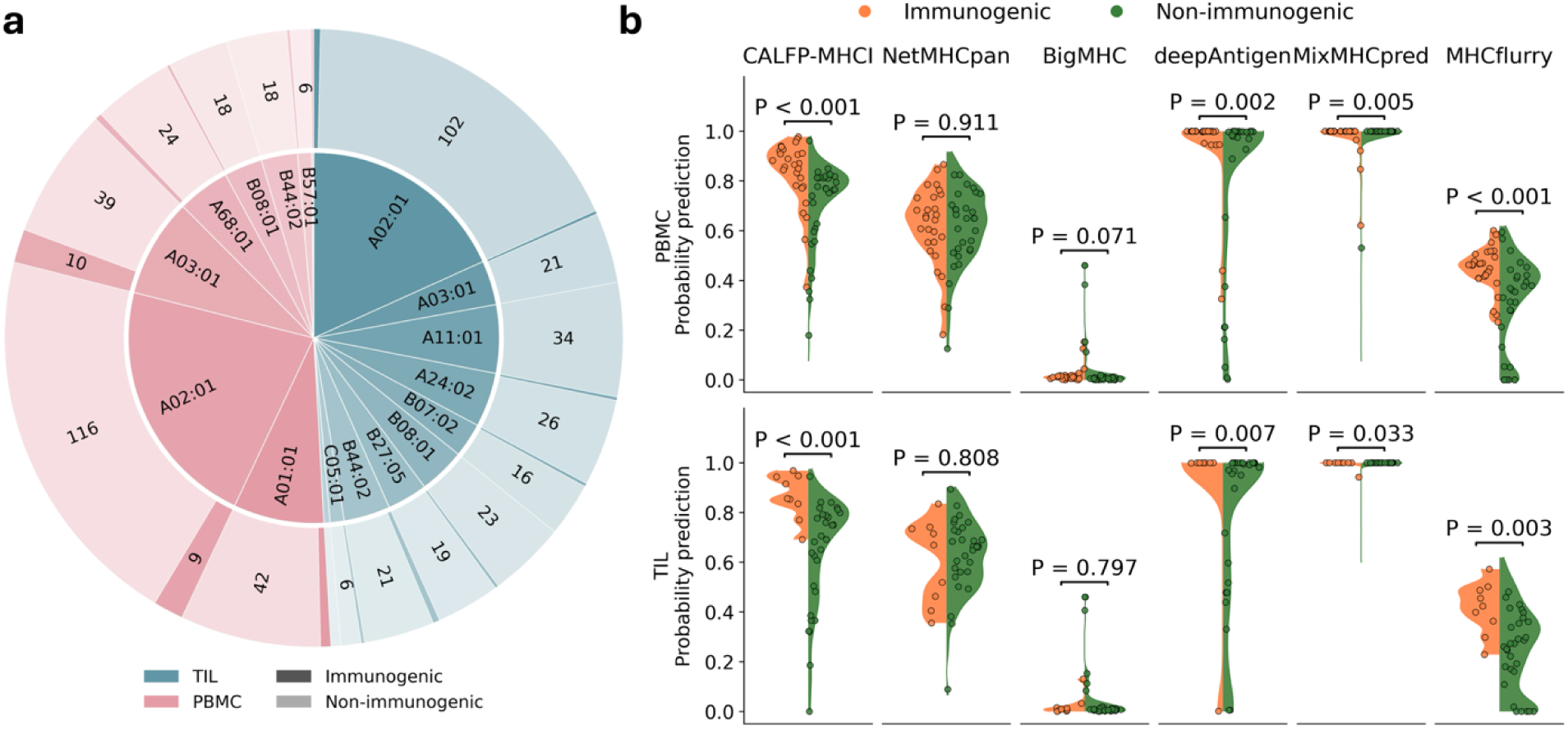
| CALFP-MHC prioritizes immunogenic neoantigens from the TESLA dataset. **a**, Distribution of immunogenic and non-immunogenic neoantigens from the TESLA dataset^23^ (570 peptide–HLA pairs after preprocessing; 36 immunogenic, 534 non-immunogenic), stratified by T cell source (TIL and PBMC) and allele identity. Immunogenicity was determined by pMHC-I multimer staining followed by flow cytometry to detect antigen-specific CD8 T cells. The inner ring shows the proportion of immunogenic (orange) and non-immunogenic (teal) neoantigens; the outer ring shows allele-specific counts, with HLA-A*02:01 representing the largest subset (n = 229 pairs). **b,** Prediction score distributions for immunogenic (orange) and non-immunogenic (teal) neoantigens across six class I predictors, stratified by T cell source: TIL (top row) and PBMC (bottom row), using the dataset shown in a. Statistical comparisons were performed using the Mann–Whitney U test (two-sided). *p < 0.05, **p < 0.01, ***p < 0.001, ns = not significant.

Overall, these results demonstrate the versatility of CALFP-MHC across diverse biological contexts, with direct relevance to neoantigen prioritization and precision immunotherapy. This robustness is sustained by consistently high predictive precision, whereas competing models show performance degradation under the same conditions.

## Discussion

In this study, we introduced CALFP-MHC, a fingerprint-based deep learning framework for peptide– MHC binding prediction that integrates residue-level chemical features, positional encoding, and supervised contrastive learning. The model consistently outperformed state-of-the-art methods^1–7,18^ across both class I and class II benchmarks, including challenging settings with severe class imbalance, rare alleles, and unseen peptides. On independent experimental datasets, MS/MS-validated MHC ligands^22^ and the TESLA neoantigen cohort^23^, CALFP-MHC maintained strong discrimination, providing a more direct assessment of performance on biologically validated data.

The fingerprint-based encoding strategy builds on prior work showing that cheminformatics descriptors can serve as effective amino acid representations^9,10^. CALFP-MHC extends this principle by embedding fingerprint representations within a hybrid convolutional transformer architecture that explicitly models positional context and long-range peptide–MHC interactions. The two-stage training strategy, using supervised contrastive pre-training to structure the latent space before fine-tuning, follows the framework of Khosla et al.^24^ and has been independently explored for MHC-II binding by ConBoTNet^11^. CALFP-MHC differs in its use of fingerprint-based inputs and pan-allelic scope across both class I and class II. The latent space reorganization observed after contrastive pre-training (Fig. 4g, k), with positive pairs contracting while negative pairs shift outward, is consistent with the representation learning dynamics reported in these prior works.

A key strength of CALFP-MHC is its interpretability. Attention and attribution analyses identified canonical anchor positions consistent with well-established structural determinants of peptide–MHC binding^15,16,20^, recovered from a model trained purely on binding labels without explicit structural supervision. Functional groups highlighted by integrated gradients attribution, including amines, hydroxyls, and aromatic rings, correspond to moieties known to mediate hydrogen bonding, charge stabilization, and hydrophobic packing at the peptide–MHC interface. We note that attention weights are not equivalent to causal feature importance and should be interpreted as indicative rather than definitive evidence; validation against crystallographic contact data would be required to confirm these assignments.

Maintaining AUC values above 0.90 and PPV of approximately 0.8 to 0.9 at negative-to-positive ratios up to 200:1 substantially exceeds competing tools under the same conditions, which we attribute partly to the balanced training strategy used during development. Future work comparing models retrained under identical balanced conditions would help disentangle the contribution of the fingerprint representation from the training strategy itself.

Nevertheless, several limitations should be considered. The fingerprint representations encode amino acid side-chain chemical properties as static descriptors and do not account for conformational flexibility, peptide folding, or binding-induced structural rearrangements. The framework also does not explicitly model upstream antigen-processing mechanisms such as proteasomal cleavage or TAP transport. The TESLA cohort^23^, which provides the most direct assessment of immunogenicity prediction, contains only 36 immunogenic peptides, providing enough power to detect large discrimination effects (AUC ≥ 0.65) but insufficient to reliably rank predictors with smaller true AUC differences. The MS/MS datasets cover a limited subset of alleles and tissue sources and may be subject to experimental biases from sample preparation and identification sensitivity.

The combination of high accuracy and biological interpretability makes CALFP-MHC particularly valuable for translational applications: identifying immunogenic peptide candidates in vaccine design, and prioritizing tumor-specific neoantigens in cancer immunotherapy, complementing approaches such as BigMHC^3^ that incorporate transfer learning from immunogenicity data. Because candidate prioritization requires both statistical accuracy and biologically meaningful evidence, the explainable framework of CALFP-MHC offers a potential advantage over conventional black-box prediction models, though prospective clinical validation remains an essential next step.

## Conclusion

In conclusion, CALFP-MHC demonstrates that integrating cheminformatics fingerprint representations with deep learning and supervised contrastive learning can improve both the accuracy and interpretability of peptide–MHC binding prediction. The framework achieved strong performance across diverse and challenging benchmark settings while identifying biologically meaningful binding motifs consistent with known structural interactions. Although further validation on larger and more diverse datasets is needed, CALFP-MHC provides a promising and interpretable approach for applications in vaccine development, neoantigen discovery, and computational immunology research.

## Methods

### Data collection and preprocessing

In this study, peptide–MHC class I and class II datasets were systematically constructed using large-scale binding predictions from established computational tools as surrogate labels. We note that these labels reflect the output of existing predictors rather than direct experimental measurements; results on this dataset therefore quantify agreement with established tools. Independent validation on experimentally measured datasets is reported separately. For class I (Fig. 7a), we employed NetMHCpan-4.1^1^ and MHCflurry-2.0^2^, generating a total of 567,816 positive peptides and 18,520,426 negatives. The data were split into training and validation sets. In the training cohort (Fig. 7b), we applied class balancing, yielding 244,974 positives and an equal number of negatives. Across loci, HLA-B was predominant with 234,433 peptides, followed by HLA-A (160,497) and HLA-C (95,018). By length, 9-mers were most frequent (209,961), while 10-mers (109,204), 8-mers (79,984), and 11-mers (90,799) were less represented. Allele-wise, HLA-A*02:01 was the most abundant with more than 400,000 peptides, followed by several HLA-B alleles such as HLA-B*07:02 and HLA-B*35:01, each exceeding 200,000 peptides. The validation cohort (Fig. 7c) comprised 322,842 positives and 18,275,454 negatives, reflecting the inherent imbalance in peptide–MHC binding predictions. Here, HLA-B again dominated (7,842,533 peptides), followed by HLA-A (6,732,303 peptides) and HLA-C (4,023,458 peptides). In terms of length, 9-mers were strongly enriched (4,808,506), alongside substantial counts of 11-mers (4,666,092), 10-mers (4,655,659), and 8-mers (4,468,037). Allele distributions demonstrated wide diversity, including common variants such as HLA-B*07:02, HLA-B*35:01, HLA-A*02:01, and HLA-A*03:01, ensuring broad and balanced coverage.

**Fig. 7.**
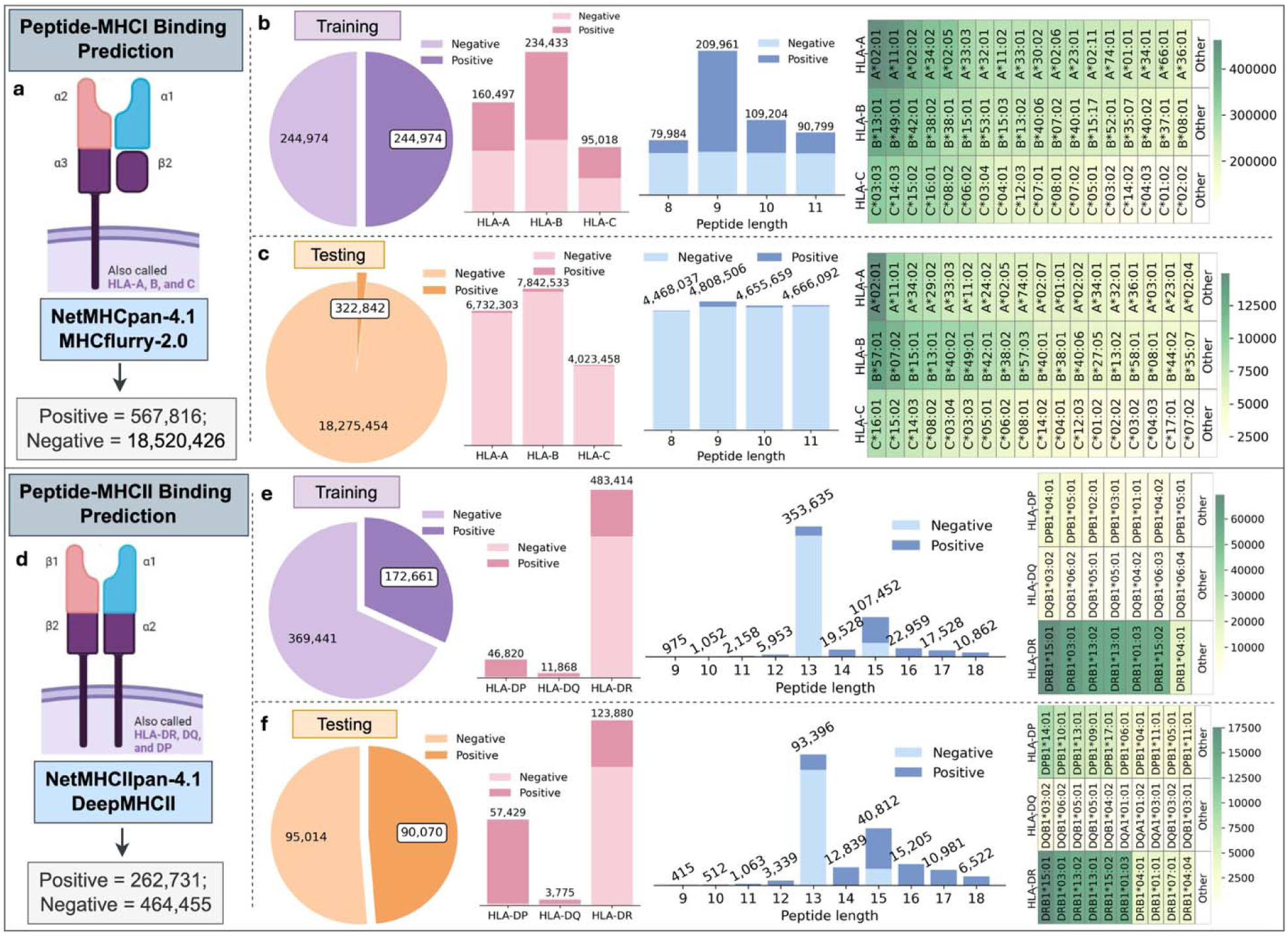
| Construction of training and testing datasets for peptide–MHC binding prediction. **a–c**, Dataset preparation for class I binding. **a,** Positive and negative peptide–MHC pairs collected from NetMHCpan-4.1^1^ and MHCflurry-2.0^2^. **b,** Training set distribution across HLA-A, HLA-B, and HLA-C alleles, stratified by peptide length (9–11 residues). **c,** Testing set distribution retaining natural class imbalance while covering diverse alleles and lengths. **d–f,** Dataset preparation for class II binding. **d,** Positive and negative peptide–MHC pairs derived from NetMHCIIpan-4.1^17^ an DeepMHCII^5^. **e,** Training set distribution across HLA-DR, HLA-DQ, and HLA-DP, stratified by peptide length (9–18 residues). **f,** Testing set distribution preserving real-world class imbalance while maintaining representative allele an length coverage.

For class II (Fig. 7d), we constructed datasets using NetMHCIIpan-4.1^17^ and DeepMHCII^5^, yielding 262,731 positives and 464,455 negatives. In the training set (Fig. 7e), there were 172,661 positives and 369,441 negatives. HLA-DR was overwhelmingly dominant with 483,414 peptides, while HLA-DP and HLA-DQ contributed only 46,820 and 11,868 peptides, respectively. By length, 15-mers were most common (107,452), followed by 16-mers (22,959), 14-mers (19,528), and 13-mers (19,528), while shorter peptides (9–12 aa) and longer ones (17–18 aa) were much rarer. The allele distribution was broad, covering multiple HLA-DRB1 variants, notably HLA-DRB1*15:01 and HLA-DRB1*04:01, each with more than 60,000 peptides. The validation set (Fig. 7f) contained 90,070 positives and 95,014 negatives. Locus-wise, HLA-DR again dominated (123,880 peptides), followed by HLA-DP (57,429) and HLA-DQ (3,775). By length, 15-mers remained the most abundant (93,396), followed by 16-mers (40,812) and 14-mers (12,839), with other lengths represented only by hundreds to a little over one thousand peptides. The allele distribution was similarly diverse, including multiple DPB1 and DRB1 variants, particularly HLA-DRB1*15:01, HLA-DRB1*04:01, and HLA-DPB1*05:01, each with high representation.

### CALFP-MHC training

#### Feature Representation

Peptides and MHC molecules were represented in a unified fingerprint-based framework that preserves both chemical detail and sequence order (Fig. 1b). Each amino acid, from either the peptide ligand or the 34-residue MHC pseudo-sequence, was encoded as a 4263-dimensional fingerprint vector^9^. The MHC allele was represented as a 34-residue pseudo-sequence, comprising the polymorphic amino acid positions located within 4.0 Å of the bound peptide across representative MHC crystal structures, as defined by Nielsen et al^13,14^. This compact encoding captures allele-specific binding specificity while remaining computationally tractable. The dimension was obtained by concatenating four widely used cheminformatics descriptors: (i) MACCS keys (166 bits), which capture predefined functional groups; (ii) ECFP4 (1024 bits) and (iii) ECFP6 (2048 bits), which describe extended circular atom neighborhoods at different radii; and (iv) RDKit molecular fingerprints (1024 bits), which encode substructure patterns and connectivity. Together, these descriptors capture complementary aspects of residue-level chemistry beyond symbolic encodings.

Residue fingerprints were stacked to form a sequence matrix, with rows representing residues and columns representing fingerprint dimensions (Fig. 1b). To incorporate positional context, each fingerprint vector at position p was combined with a sinusoidal positional encoding via element-wise addition, following the approach of Vaswani et al^12^. This additive encoding preserves the magnitude of fingerprint features while embedding residue position information independently across dimensions. This step differentiates sequences with identical amino acid compositions but different residue orders, embedding sequence order directly in the fingerprint space.

Following positional adjustment, the sequence matrices were normalized by column-wise summation and division by the maximum value, scaling all features into the [0,1] range. This produced compact sequence-level vectors for both peptides and MHC pseudo-sequences. The two vectors were then concatenated to form a joint interaction fingerprint that integrates residue-level chemistry with positional information, providing the standardized input to the downstream deep learning model for peptide–MHC binding.

#### Model Architecture

Our model is designed to capture the complex dependencies between peptides and MHC molecules through a structured deep learning framework (Fig. 1c-d). The architecture consists of four main components: (i) a fingerprint-based sequence encoder, (ii) a backbone network with bottleneck transformer blocks, (iii) a supervised contrastive pre-training stage, and (iv) a fine-tuning stage for binding affinity prediction and binding core identification.

#### Sequence encoder

Each peptide of length L and each 34-residue MHC pseudo-sequence were first represented as residue-level fingerprint vectors of dimension *d* = 4,263, yielding matrices 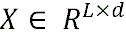 and 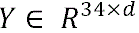 To align the MHC sequence representation with the peptide binding core of length *k*, we applied a learnable linear projection:

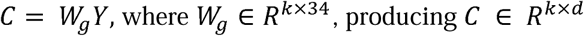

Here, each row of C acts as a kernel conditioned on the MHC pseudo-sequence.

For each sliding window of length k over the peptide, element-wise interactions were computed between the peptide fragment and the kernel generated from the MHC pseudo-sequence (Fig. 1c):

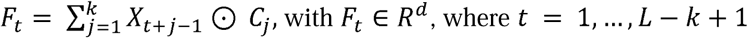

Where ⊙ denotes element-wise multiplication. Concatenating all *F_t_* produces an interaction matrix 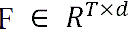, with *T* = *L* – *k* + 1, which serves as the input to the backbone encoder.

This combined fingerprint encodes residue-level chemistry and position information for both peptide and MHC.

#### Backbone network with bottlenecked transformer architecture

The backbone uses a bottleneck transformer, which combines convolutional filters, residual connections, and self-attention. The bottleneck reduces dimensionality to lower computational cost while preserving long-range dependencies in peptide–MHC interactions.

Dimensionality reduction. The interaction features are first projected into a lower-dimensional space:

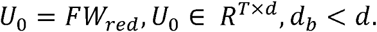

Multi-head self-attention (MHSA). Queries, keys, and values are computed as:

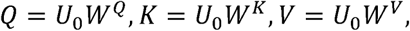

with 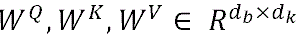, and 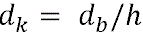 for *h* attention heads. The attention update is:

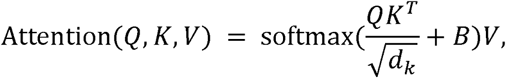

Where 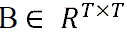 encodes relative positional bias. Outputs from all heads are concatenated:

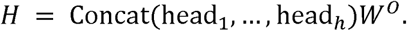

Residual connections and feed-forward update.

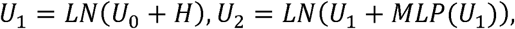

Expansion and convolution refinement.

The reduced representation is projected back to the original dimension, merged with the input, and passed through residual convolutional blocks:

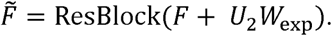

Finally, average pooling across sequence positions produces a global representation:

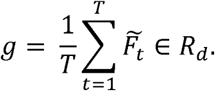

#### Supervised contrastive pre-training

To impose a structured latent space, peptide–MHC pairs were labeled as binder or non-binder. The pooled representation g is mapped through a two-layer projection head:

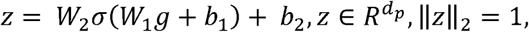

with ReLU non-linearity and normalization on the unit sphere. The supervised contrastive loss is:

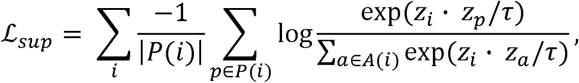

where *p*(*i*) is the set of positives with the same class label, *A*(*i*), the remaining samples, and *τ* a temperature parameter. This loss clusters embeddings of binders together while separating them from non-binders (Fig. 1d), yielding a discriminative latent space.

#### Fine-tuning for binding affinity prediction

During fine-tuning, the projection head is replaced with a binary classification layer:

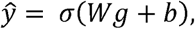

where 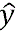 represents the predicted probability of binding. The model is optimized using binary cross-entropy loss (Fig. 1d – Stage 2):

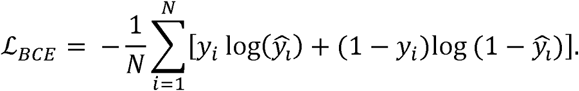

This setup enables the encoder to first learn a discriminative latent structure and then specialize for accurate binding prediction.

#### Training hyperparameters

CALFP-MHC was implemented with the following architecture. Each residue was encoded as a 4,263-dimensional fingerprint vector; the interaction layer processed peptide–MHC pairs through convolutional filters (3,200 channels, kernel size 9, stride 1) to produce a 256-dimensional joint feature representation. High-level features were refined by a bottleneck transformer (2 blocks, 9 attention heads, bottleneck dimension 256) combined with ResNet blocks (1-D depthwise convolutions, kernel size 9, 3,200 channels, GLU activation, identity skip connections). The contrastive projection head used two fully connected layers (800 → 64) with SiLU activation and L2 normalisation. The EL and BA output heads each comprised a single linear layer from 64-dimensional embeddings, with softmax and sigmoid activations respectively. Dropout of 0.2 was applied after each attention and convolutional block.

Training proceeded in two stages using the Adam optimizer with weight decay 1×10□□. In Stage 1, the full network was trained with supervised contrastive loss (τ = 0.07). In Stage 2, the encoder was frozen and the dual output heads were fine-tuned independently (batch size = 256, learning rate = 1×10□□, 100 epochs, early stopping patience 10). All experiments were run on NVIDIA A100 GPU with CUDA 11.4.

#### Binding score identification

Attention scores from the transformer blocks are aggregated across positions to estimate the importance of peptide residues (Fig. 1d – Stage 2). Sub-sequences with the highest cumulative attention are reported as candidate binding cores, providing mechanistic interpretability in addition to predictive performance.

### Comparison of CALFP-MHC with other methods on the independent test set

All datasets were systematically curated to include a comprehensive range of HLA alleles across both MHCI and MHCII molecules. For each allele, the number of positive and negative peptide–HLA pairs was recorded (Table S1 and S2). In the case of MHCI, a total of 112 alleles were considered and further grouped into three classes, HLA-A, HLA-B, and HLA-C, with HLA-B contributing the largest number of pairs (6,724,864 pairs). For MHCII, 53 alleles were analyzed and classified into two major groups, HLA-DR and HLA-DP, where HLA-DR accounted for the highest representation (733,860 pairs). The performance of the models across these allele groups is presented in Fig. 2 and Fig. 3.

To benchmark the proposed CALFP framework, we evaluated both MHCI and MHCII models against widely used predictive tools. For MHCI, comparisons were made with NetMHCpan, MixMHCpred^4^, MHCflurry, deepAntigen, and BigMHC, while for MHCII, benchmarking included NetMHCIIpan^17^, MixMHC2pred^6^, DeepSeqPanII^7^, DeepMHCII^5^, and Tlimmuno2^18^. Two complementary evaluation strategies were employed. First, datasets were partitioned according to the ratio of positive and negative peptide–HLA pairs, which resulted in varying proportions of seen and unseen peptides depending on the training sets of individual tools (Tables S3). Second, an allele-based evaluation was performed, in which MHCI peptides were grouped into HLA-A, HLA-B, and HLA-C, and MHCII peptides were grouped into HLA-DR and HLA-DP. The distribution of seen and unseen samples across tools under this scheme is summarized in Tables S4.

### Application of validated neoantigen and experimental data

In addition to the training and testing datasets, we collected independent non-overlapping datasets to further evaluate the performance of CALFP-MHC. These datasets included validated neoantigen experiments, mass spectrometry (MS/MS) assays for MHCI, and both IC50 binding assays and MS/MS data for MHCII. For MHCI, we curated the TESLA dataset^23^ comprising 608 peptide–HLA pairs with 37 positive and 571 negative cases. After preprocessing, which involved the removal of duplicates and overlaps with the training and testing sets, 570 pairs remained with 36 positive and 534 negative, spanning 13 HLA alleles, with HLA-A*02:01 contributing the largest subset of 229 pairs. MS/MS-derived data for MHCI were obtained^22^, initially including 3,472 positive and 27,632 negative cases. After data cleaning, we retained 2,827 positive and 800 negative, covering 16 HLA alleles, and no allele showed a dominant imbalance in the number of pairs. For MHCII, we assembled two complementary datasets^17^, and after preprocessing, the combined MHCII dataset contained 520 pairs, with 306 positive and 214 negative, of which 390 pairs came from MS/MS assays and 130 pairs from IC50 assays. Among these, HLA-DQA1*0301 accounted for the largest number of pairs. Detailed statistics of all collected datasets before and after preprocessing are summarized in Table S6.

### Statistical analysis

Score distributions between positive and negative peptides were compared using the Mann-Whitney U test (two-sided); p-values are indicated above each distribution. Significance thresholds: *p < 0.05, **p < 0.01, ***p < 0.001, ns = not significant. No multiple testing correction was applied as each comparison was pre-specified. AUC values and 95% bootstrap confidence intervals were computed using 1,000 bootstrap resamples. Pairwise AUC comparisons on the TESLA dataset were performed using DeLong’s method. Statistical analyses were conducted using Python scikit-learn and scipy.stats.

## Data availability

All data were collected from publicly available sources: MHCflurry-2.0^2^, NetMHCpan-4.1^1^, NetMHCIIpan-4.1^17^, DeepMHCII^5^, TESLA^23^

## Code availability

The source code used in this study is openly available at GitHub: https://github.com/ddiem-ri-4D/CALFP-MHC. Installation instructions and usage examples are provided in the README file.

## Declaration of generative AI use

During the preparation of this manuscript, the authors used ChatGPT (OpenAI) to assist with improving the language, grammar, readability, and clarity of the manuscript. The authors remain fully responsible for the content, interpretation of results, scientific accuracy, and conclusions presented in this work. No generative AI tool was used to generate, analyze, or interpret the underlying scientific data.

## Acknowledgements

This research is funded by NexCalibur Therapeutics under grant number NCT01.

MDP holds shares in NexCalibur Therapeutics, the company that has provided funding for the research presented in this publication.

