## Supplementary Info for "CALFP-MHC: Interpretable Pan-Allelic Prediction of Peptide-MHC Binding and Presentation Using Chemically Grounded Fingerprints and Contrastive Learning"

### Supplementary Methods

#### CALFP-MHCI outperformed existing tools in predicting the binding between HLAI-peptide

### Supplementary Tables

Table S1:

| No. | Allele | n pairs | pos | neg | seen  peps | unsee  peps | No. | Allele | n  pairs | pos | neg | seen  peps | unsee  peps |
| --- | --- | --- | --- | --- | --- | --- | --- | --- | --- | --- | --- | --- | --- |
| 1 | A*01:01 | 226305 | 975 | 225330 | 540 | 225765 | 57 | B*39:06 | 14840 | 495 | 14345 | 563 | 14277 |
| 2 | A*02:01 | 447621 | 4484 | 443137 | 2864 | 444757 | 58 | B*39:10 | 45 | 5 | 40 | 6 | 39 |
| 3 | A*02:03 | 121623 | 307 | 121316 | 3180 | 118443 | 59 | B*39:24 | 2812 | 259 | 2553 | 284 | 2528 |
| 4 | A*02:06 | 261344 | 2033 | 259311 | 8802 | 252542 | 60 | B*40:01 | 224086 | 4783 | 219303 | 11278 | 212808 |
| 5 | A*02:07 | 114100 | 483 | 113617 | 2152 | 111948 | 61 | B*40:02 | 192072 | 8720 | 183352 | 13110 | 178962 |
| 6 | A*11:01 | 439272 | 2722 | 436550 | 10105 | 429167 | 62 | B*40:06 | 239442 | 2259 | 237183 | 12399 | 227043 |
| 7 | A*11:02 | 263808 | 278 | 263530 | 7871 | 255937 | 63 | B*41:01 | 14082 | 451 | 13631 | 559 | 13523 |
| 8 | A*24:02 | 129484 | 1139 | 128345 | 891 | 128593 | 64 | B*41:02 | 157 | 17 | 140 | 19 | 138 |
| 9 | A*24:07 | 142536 | 124 | 142412 | 4185 | 138351 | 65 | B*41:03 | 664 | 80 | 584 | 90 | 574 |
| 10 | A*26:01 | 151161 | 650 | 150511 | 3766 | 147395 | 66 | B*41:04 | 298 | 38 | 260 | 42 | 256 |
| 11 | A*33:03 | 287427 | 295 | 287132 | 7700 | 279727 | 67 | B*41:06 | 196 | 16 | 180 | 18 | 178 |
| 12 | A*02:02 | 380552 | 3267 | 377285 | 16866 | 363686 | 68 | B*42:01 | 342674 | 3230 | 339444 | 17490 | 325184 |
| 13 | A*02:04 | 95267 | 3412 | 91855 | 6224 | 89043 | 69 | B*44:02 | 111880 | 4615 | 107265 | 5374 | 106506 |
| 14 | A*02:05 | 343213 | 3250 | 339963 | 15943 | 327270 | 70 | B*44:03 | 105399 | 4453 | 100946 | 5334 | 100065 |
| 15 | A*02:11 | 248312 | 2066 | 246246 | 11761 | 236551 | 71 | B*44:05 | 6177 | 4 | 6173 | 4 | 6173 |
| 16 | A*03:01 | 143781 | 4600 | 139181 | 5407 | 138374 | 72 | B*44:08 | 626 | 47 | 579 | 50 | 576 |
| 17 | A*23:01 | 263689 | 2631 | 261058 | 12489 | 251200 | 73 | B*44:27 | 1536 | 178 | 1358 | 195 | 1341 |
| 18 | A*24:13 | 3101 | 228 | 2873 | 273 | 2828 | 74 | B*45:01 | 162659 | 2135 | 160524 | 6933 | 155726 |
| 19 | A*25:01 | 57566 | 1064 | 56502 | 3248 | 54318 | 75 | B*46:01 | 98255 | 2441 | 95814 | 5584 | 92671 |
| 20 | A*29:02 | 154030 | 7002 | 147028 | 8774 | 145256 | 76 | B*47:01 | 366 | 26 | 340 | 27 | 339 |
| 21 | A*30:01 | 128891 | 1386 | 127505 | 5570 | 123321 | 77 | B*49:01 | 387866 | 3874 | 383992 | 19670 | 368196 |
| 22 | A*30:02 | 267497 | 2342 | 265155 | 12152 | 255345 | 78 | B*50:01 | 71517 | 1110 | 70407 | 3569 | 67948 |
| 23 | A*31:01 | 32866 | 1802 | 31064 | 2613 | 30253 | 79 | B*51:01 | 121994 | 3943 | 118051 | 5329 | 116665 |
| 24 | A*32:01 | 272559 | 2683 | 269876 | 11566 | 260993 | 80 | B*51:02 | 67 | 7 | 60 | 8 | 59 |
| 25 | A*68:01 | 106960 | 1846 | 105114 | 5848 | 101112 | 81 | B*51:08 | 14043 | 616 | 13427 | 795 | 13248 |
| 26 | A*68:02 | 51568 | 1922 | 49646 | 3285 | 48283 | 82 | B*52:01 | 218140 | 1726 | 216414 | 10955 | 207185 |
| 27 | B*07:02 | 235210 | 11058 | 224152 | 13462 | 221748 | 83 | B*53:01 | 247639 | 2161 | 245478 | 11074 | 236565 |
| 28 | B*08:01 | 180807 | 4857 | 175950 | 6235 | 174572 | 84 | B*54:01 | 40514 | 1261 | 39253 | 2763 | 37751 |
| 29 | B*13:02 | 246166 | 2225 | 243941 | 12848 | 233318 | 85 | B*55:01 | 149083 | 1454 | 147629 | 7392 | 141691 |
| 30 | B*14:02 | 168120 | 1523 | 166597 | 6332 | 161788 | 86 | B*55:02 | 148577 | 1430 | 147147 | 8287 | 140290 |
| 31 | B*15:01 | 273654 | 8147 | 265507 | 14659 | 258995 | 87 | B*56:01 | 144361 | 1947 | 142414 | 7640 | 136721 |
| 32 | B*15:17 | 222370 | 1741 | 220629 | 9326 | 213044 | 88 | B*57:01 | 177068 | 13061 | 164007 | 14545 | 162523 |
| 33 | B*15:18 | 1427 | 5 | 1422 | 5 | 1422 | 89 | B*57:03 | 171680 | 5589 | 166091 | 11721 | 159959 |
| 34 | B*18:01 | 161666 | 2622 | 159044 | 7012 | 154654 | 90 | B*58:01 | 142617 | 4545 | 138072 | 8519 | 134098 |
| 35 | B*18:03 | 2289 | 212 | 2077 | 251 | 2038 | 91 | B*58:02 | 107485 | 994 | 106491 | 5789 | 101696 |
| 36 | B*27:01 | 43024 | 3653 | 39371 | 4138 | 38886 | 92 | B*73:01 | 2250 | 175 | 2075 | 213 | 2037 |
| 37 | B*27:02 | 45066 | 2853 | 42213 | 3218 | 41848 | 93 | C*01:02 | 112011 | 2312 | 109699 | 6007 | 106004 |
| 38 | B*27:03 | 10209 | 887 | 9322 | 1033 | 9176 | 94 | C*02:02 | 108882 | 2385 | 106497 | 5368 | 103514 |
| 39 | B*27:04 | 16752 | 1222 | 15530 | 1458 | 15294 | 95 | C*03:02 | 127268 | 1206 | 126062 | 7126 | 120142 |
| 40 | B*27:05 | 89805 | 5052 | 84753 | 6594 | 83211 | 96 | C*03:03 | 329535 | 3509 | 326026 | 14812 | 314723 |
| 41 | B*27:06 | 16232 | 1139 | 15093 | 1415 | 14817 | 97 | C*03:04 | 218990 | 3580 | 215410 | 10745 | 208245 |
| 42 | B*27:07 | 24230 | 1958 | 22272 | 2302 | 21928 | 98 | C*04:01 | 205198 | 2871 | 202327 | 6984 | 198214 |
| 43 | B*27:08 | 21165 | 1895 | 19270 | 2196 | 18969 | 99 | C*04:03 | 113681 | 1069 | 112612 | 6261 | 107420 |
| 44 | B*27:09 | 54871 | 4292 | 50579 | 5018 | 49853 | 100 | C*05:01 | 168001 | 4819 | 163182 | 8048 | 159953 |
| 45 | B*27:10 | 130 | 10 | 120 | 13 | 117 | 101 | C*06:02 | 266223 | 3117 | 263106 | 9624 | 256599 |
| 46 | B*35:01 | 76691 | 2358 | 74333 | 3351 | 73340 | 102 | C*07:01 | 201499 | 1160 | 200339 | 4391 | 197108 |
| 47 | B*35:02 | 27074 | 34 | 27040 | 39 | 27035 | 103 | C*07:02 | 183085 | 1904 | 181181 | 5226 | 177859 |
| 48 | B*35:03 | 133311 | 1496 | 131815 | 6076 | 127235 | 104 | C*07:04 | 106417 | 794 | 105623 | 5287 | 101130 |
| 49 | B*35:04 | 64 | 5 | 59 | 6 | 58 | 105 | C*08:01 | 198933 | 1813 | 197120 | 10761 | 188172 |
| 50 | B*35:06 | 65 | 5 | 60 | 5 | 60 | 106 | C*08:02 | 275106 | 5061 | 270045 | 12316 | 262790 |
| 51 | B*35:07 | 201995 | 1819 | 200176 | 10777 | 191218 | 107 | C*12:02 | 46276 | 1378 | 44898 | 3018 | 43258 |
| 52 | B*35:08 | 2275 | 196 | 2079 | 222 | 2053 | 108 | C*12:03 | 202675 | 2855 | 199820 | 8878 | 193797 |
| 53 | B*37:01 | 180846 | 1659 | 179187 | 7528 | 173318 | 109 | C*14:02 | 127157 | 3122 | 124035 | 7019 | 120138 |
| 54 | B*38:01 | 276447 | 2471 | 273976 | 13416 | 263031 | 110 | C*14:03 | 306912 | 2817 | 304095 | 16030 | 290882 |
| 55 | B*38:02 | 320200 | 2892 | 317308 | 16431 | 303769 | 111 | C*15:02 | 296874 | 4149 | 292725 | 15269 | 281605 |
| 56 | B*39:01 | 29536 | 971 | 28565 | 1152 | 28384 | 112 | C*16:01 | 296159 | 4962 | 291197 | 14608 | 281551 |

Table S2:

| No. | Allele | n pairs | pos | neg | seen  peps | unseen  peps | No. | Allele | n  pairs | pos | neg | seen  peps | unseen  peps |
| --- | --- | --- | --- | --- | --- | --- | --- | --- | --- | --- | --- | --- | --- |
| 1 | DRB1*03:01 | 15409 | 1468 | 13941 | 593 | 14816 | 28 | DRB1*04:08 | 34 | 33 | 1 | 0 | 34 |
| 2 | DRB1*04:05 | 1205 | 1121 | 84 | 418 | 787 | 29 | DRB1*07:01 | 3802 | 1892 | 1910 | 717 | 3085 |
| 3 | DRB1*09:01 | 629 | 510 | 119 | 533 | 96 | 30 | DRB1*08:01 | 1168 | 1114 | 54 | 192 | 976 |
| 4 | DRB1*10:01 | 727 | 709 | 18 | 28 | 699 | 31 | DRB1*08:02 | 428 | 316 | 112 | 397 | 31 |
| 5 | DRB1*15:02 | 14109 | 294 | 13815 | 0 | 14109 | 32 | DRB1*08:03 | 393 | 58 | 335 | 6 | 387 |
| 6 | DPB1*01:01 | 125 | 100 | 25 | 124 | 1 | 33 | DRB1*11:01 | 1448 | 1110 | 338 | 615 | 833 |
| 7 | DPB1*02:01 | 243 | 202 | 41 | 202 | 41 | 34 | DRB1*11:02 | 7 | 2 | 5 | 0 | 7 |
| 8 | DPB1*03:01 | 144 | 109 | 35 | 126 | 18 | 35 | DRB1*11:03 | 121 | 113 | 8 | 0 | 121 |
| 9 | DPB1*04:01 | 376 | 206 | 170 | 190 | 186 | 36 | DRB1*11:04 | 28 | 16 | 12 | 0 | 28 |
| 10 | DPB1*04:02 | 319 | 188 | 131 | 181 | 138 | 37 | DRB1*12:01 | 732 | 642 | 90 | 199 | 533 |
| 11 | DPB1*05:01 | 191 | 125 | 66 | 131 | 60 | 38 | DRB1*13:01 | 14443 | 721 | 13722 | 194 | 14249 |
| 12 | DPB1*09:01 | 32 | 15 | 17 | 0 | 32 | 39 | DRB1*13:02 | 14460 | 651 | 13809 | 452 | 14008 |
| 13 | DPB1*14:01 | 214 | 144 | 70 | 198 | 16 | 40 | DRB1*14:01 | 114 | 113 | 1 | 0 | 114 |
| 14 | DPB1*15:01 | 34 | 10 | 24 | 2 | 32 | 41 | DRB1*14:02 | 7 | 4 | 3 | 0 | 7 |
| 15 | DPB1*20:01 | 175 | 144 | 31 | 175 | 0 | 42 | DRB1*14:05 | 1949 | 333 | 1616 | 30 | 1919 |
| 16 | DRB1*01:01 | 4312 | 3994 | 318 | 1646 | 2666 | 43 | DRB1*14:54 | 360 | 344 | 16 | 173 | 187 |
| 17 | DRB1*01:02 | 811 | 807 | 4 | 1 | 810 | 44 | DRB1*15:01 | 17562 | 1899 | 15663 | 542 | 17020 |
| 18 | DRB1*01:03 | 14015 | 303 | 13712 | 11 | 14004 | 45 | DRB1*15:03 | 7 | 1 | 6 | 1 | 6 |
| 19 | DRB1*03:02 | 17 | 5 | 12 | 6 | 11 | 46 | DRB1*16:01 | 71 | 66 | 5 | 0 | 71 |
| 20 | DRB1*03:03 | 7 | 5 | 2 | 1 | 6 | 47 | DRB1*16:02 | 27 | 22 | 5 | 24 | 3 |
| 21 | DRB1*03:05 | 2 | 1 | 1 | 0 | 2 | 48 | DRB3*01:01 | 1019 | 810 | 209 | 520 | 499 |
| 22 | DRB1*04:01 | 4520 | 4085 | 435 | 589 | 3931 | 49 | DRB3*02:02 | 1077 | 920 | 157 | 332 | 745 |
| 23 | DRB1*04:02 | 1320 | 1279 | 41 | 23 | 1297 | 50 | DRB3*03:01 | 185 | 171 | 14 | 156 | 29 |
| 24 | DRB1*04:03 | 13 | 7 | 6 | 0 | 13 | 51 | DRB4*01:01 | 587 | 467 | 120 | 404 | 183 |
| 25 | DRB1*04:04 | 2969 | 1147 | 1822 | 526 | 2443 | 52 | DRB4*01:03 | 828 | 800 | 28 | 185 | 643 |
| 26 | DRB1*04:06 | 7 | 6 | 1 | 4 | 3 | 53 | DRB5*01:01 | 1341 | 1174 | 167 | 588 | 753 |
| 27 | DRB1*04:07 | 40 | 37 | 3 | 0 | 40 |  |  |  |  |  |  |  |

Table S3:

|  |  | neg=1.pos | | | | | neg=10.pos | | | | |
| --- | --- | --- | --- | --- | --- | --- | --- | --- | --- | --- | --- |
|  |  | Overall | Seen | Unseen peptides | Unseen MHC | Unseen Peptide + MHC | Overall | Seen | Unseen peptides | Unseen MHC | Unseen Peptide + MHC |
| MHCI | CALFP-MHCI | 155738 | 36002 | 119483 | 1 | 252 | 856559 | 51163 | 802987 | 1 | 2408 |
|  | NetMHCpan |  |  |  |  |  |  |  |  |  |  |
|  | MixMHCpred | 645684 | 257536 | 374665 | 7771 | 5712 | 3551262 | 292029 | 3223296 | 7787 | 28150 |
|  | MHCflurry | 645684 | 0 | 0 | 16726 | 628958 | 3551262 | 0 | 0 | 109896 | 3441366 |
|  | deepAntigen | 645684 | 186169 | 429402 | 6627 | 23486 | 3551262 | 222382 | 3233685 | 6659 | 88536 |
|  | BigMHC | 645684 | 575849 | 68861 | 4 | 970 | 3551262 | 3027068 | 514550 | 34 | 9610 |
| MHCII | CALFP-MHCII | 180140 | 9787 | 105086 | 3506 | 61761 | 104511 | 2961 | 91165 | 1334 | 9051 |
|  | NetMHCIIpan | 180140 | 8084 | 106789 | 2851 | 62416 | 104511 | 2493 | 91633 | 1100 | 9285 |
|  | MixMHC2pred | 180140 | 114 | 176022 | 9 | 4521 | 104511 | 17 | 103249 | 1 | 1255 |
|  | DeepSeqPanII | 180140 | 1956 | 22129 | 11340 | 144715 | 104511 | 875 | 2808 | 3420 | 97408 |
|  | DeepMHCII | 180140 | 9787 | 105086 | 3506 | 61761 | 104511 | 2961 | 91165 | 1334 | 9051 |
|  | Tlimmuno2 | 180140 | 8154 | 107193 | 2768 | 62025 | 104511 | 2521 | 91671 | 1089 | 9230 |
|  |  | neg=20.pos | | | | | neg=50.pos | | | | |
|  |  | Overall | Seen | Unseen peptides | Unseen MHC | Unseen Peptide + MHC | Overall | Seen | Unseen peptides | Unseen MHC | Unseen Peptide + MHC |
| MHCI | CALFP-MHCI | 1635249 | 68156 | 1562350 | 1 | 4742 | 3971319 | 118674 | 3840943 | 3 | 11699 |
|  | NetMHCpan |  |  |  |  |  |  |  |  |  |  |
|  | MixMHCpred | 6779682 | 330754 | 6388018 | 7808 | 53102 | 16464942 | 446144 | 15883010 | 7857 | 127931 |
|  | MHCflurry | 6779682 | 0 | 0 | 213367 | 6566315 | 16464942 | 0 | 0 | 523230 | 15941712 |
|  | deepAntigen | 6779682 | 262972 | 6349680 | 6695 | 160335 | 16464942 | 384177 | 15696505 | 6787 | 377473 |
|  | BigMHC | 6779682 | 5751694 | 1008563 | 56 | 19369 | 16464942 | 13925485 | 2490823 | 147 | 48487 |
| MHCII | CALFP-MHCII | 180140 | 9787 | 105086 | 3506 | 61761 | 104511 | 2961 | 91165 | 1334 | 9051 |
|  | NetMHCIIpan | 180140 | 8084 | 106789 | 2851 | 62416 | 104511 | 2493 | 91633 | 1100 | 9285 |
|  | MixMHC2pred | 180140 | 114 | 176022 | 9 | 4521 | 104511 | 17 | 103249 | 1 | 1255 |
|  | DeepSeqPanII | 180140 | 1956 | 22129 | 11340 | 144715 | 104511 | 875 | 2808 | 3420 | 97408 |
|  | DeepMHCII | 180140 | 9787 | 105086 | 3506 | 61761 | 104511 | 2961 | 91165 | 1334 | 9051 |
|  | Tlimmuno2 | 180140 | 8154 | 107193 | 2768 | 62025 | 104511 | 2521 | 91671 | 1089 | 9230 |
|  |  | neg=100.pos | | | | | neg=200.pos | | | | |
|  |  | Overall | Seen | Unseen peptides | Unseen MHC | Unseen Peptide + MHC | Overall | Seen | Unseen peptides | Unseen MHC | Unseen Peptide + MHC |
| MHCI | CALFP-MHCI | 7864769 | 202250 | 7639011 | 3 | 23505 | 15651669 | 370531 | 15233514 | 5 | 47619 |
|  | NetMHCpan |  |  |  |  |  |  |  |  |  |  |
|  | MixMHCpred | 18598294 | 471575 | 17974130 | 7864 | 144725 | 18598294 | 471575 | 17974130 | 7864 | 144725 |
|  | MHCflurry | 18598294 | 0 | 0 | 591219 | 18007075 | 18598294 | 0 | 0 | 591219 | 18007075 |
|  | deepAntigen | 18598294 | 410597 | 17755705 | 6808 | 425184 | 18598294 | 410597 | 17755705 | 6808 | 425184 |
|  | BigMHC | 18598294 | 15724367 | 2818767 | 167 | 54993 | 18598294 | 15724367 | 2818767 | 167 | 54993 |
| MHCII | CALFP-MHCII | 180140 | 9787 | 105086 | 3506 | 61761 | 104511 | 2961 | 91165 | 1334 | 9051 |
|  | NetMHCIIpan | 180140 | 8084 | 106789 | 2851 | 62416 | 104511 | 2493 | 91633 | 1100 | 9285 |
|  | MixMHC2pred | 180140 | 114 | 176022 | 9 | 4521 | 104511 | 17 | 103249 | 1 | 1255 |
|  | DeepSeqPanII | 180140 | 1956 | 22129 | 11340 | 144715 | 104511 | 875 | 2808 | 3420 | 97408 |
|  | DeepMHCII | 180140 | 9787 | 105086 | 3506 | 61761 | 104511 | 2961 | 91165 | 1334 | 9051 |
|  | Tlimmuno2 | 180140 | 8154 | 107193 | 2768 | 62025 | 104511 | 2521 | 91671 | 1089 | 9230 |

Table S4:

|  |  | HLA-A | | | | HLA-B | | | | HLA-C | | | |
| --- | --- | --- | --- | --- | --- | --- | --- | --- | --- | --- | --- | --- | --- |
|  |  | Seen peptides | Unseen peptides | Seen MHC | Unseen MHC | Seen peptides | Unseen peptides | Seen MHC | Unseen MHC | Seen peptides | Unseen peptides | Seen MHC | Unseen MHC |
| MHCI | CALFP-MHCI | 4782699 | 664899 | 42 | 2 | 5646413 | 600021 | 85 | 4 | 2830270 | 481940 | 22 | 4 |
|  | NetMHCpan | 707815 | 4739783 | 37 | 7 | 1191487 | 5054947 | 83 | 6 | 470360 | 2841850 | 18 | 8 |
|  | MixMHCpred | 99824 | 5347774 | 34 | 10 | 125511 | 6120923 | 59 | 30 | 66330 | 3245880 | 23 | 3 |
|  | MHCflurry | 131841 | 5315757 | 43 | 1 | 162964 | 6083470 | 87 | 2 | 78085 | 3234125 | 24 | 2 |
|  | deepAntigen | 93802 | 5353796 | 31 | 13 | 104528 | 6141906 | 40 | 49 | 58641 | 3253569 | 21 | 5 |
|  | BigMHC | 214429 | 5233169 | 42 | 2 | 259840 | 5986594 | 85 | 4 | 135245 | 3176965 | 22 | 4 |
|  |  | HLA-DP | | | | HLA-DR | | | |  |  |  |  |
|  |  | Seen peptides | Unseen peptides | Seen MHC | Unseen MHC | Seen peptides | Unseen peptides | Seen MHC | Unseen MHC |  |  |  |  |
| MHCII | CALFP-MHCII | 9787 | 105086 | 3506 | 61761 | 104511 | 2961 | 91165 | 1334 |  |  |  |  |
|  | NetMHCIIpan | 8084 | 106789 | 2851 | 62416 | 104511 | 2493 | 91633 | 1100 |  |  |  |  |
|  | MixMHC2pred | 114 | 176022 | 9 | 4521 | 104511 | 17 | 103249 | 1 |  |  |  |  |
|  | DeepSeqPanII | 1956 | 22129 | 11340 | 144715 | 104511 | 875 | 2808 | 3420 |  |  |  |  |
|  | DeepMHCII | 9787 | 105086 | 3506 | 61761 | 104511 | 2961 | 91165 | 1334 |  |  |  |  |
|  | Tlimmuno2 | 8154 | 107193 | 2768 | 62025 | 104511 | 2521 | 91671 | 1089 |  |  |  |  |

### Supplementary Figures

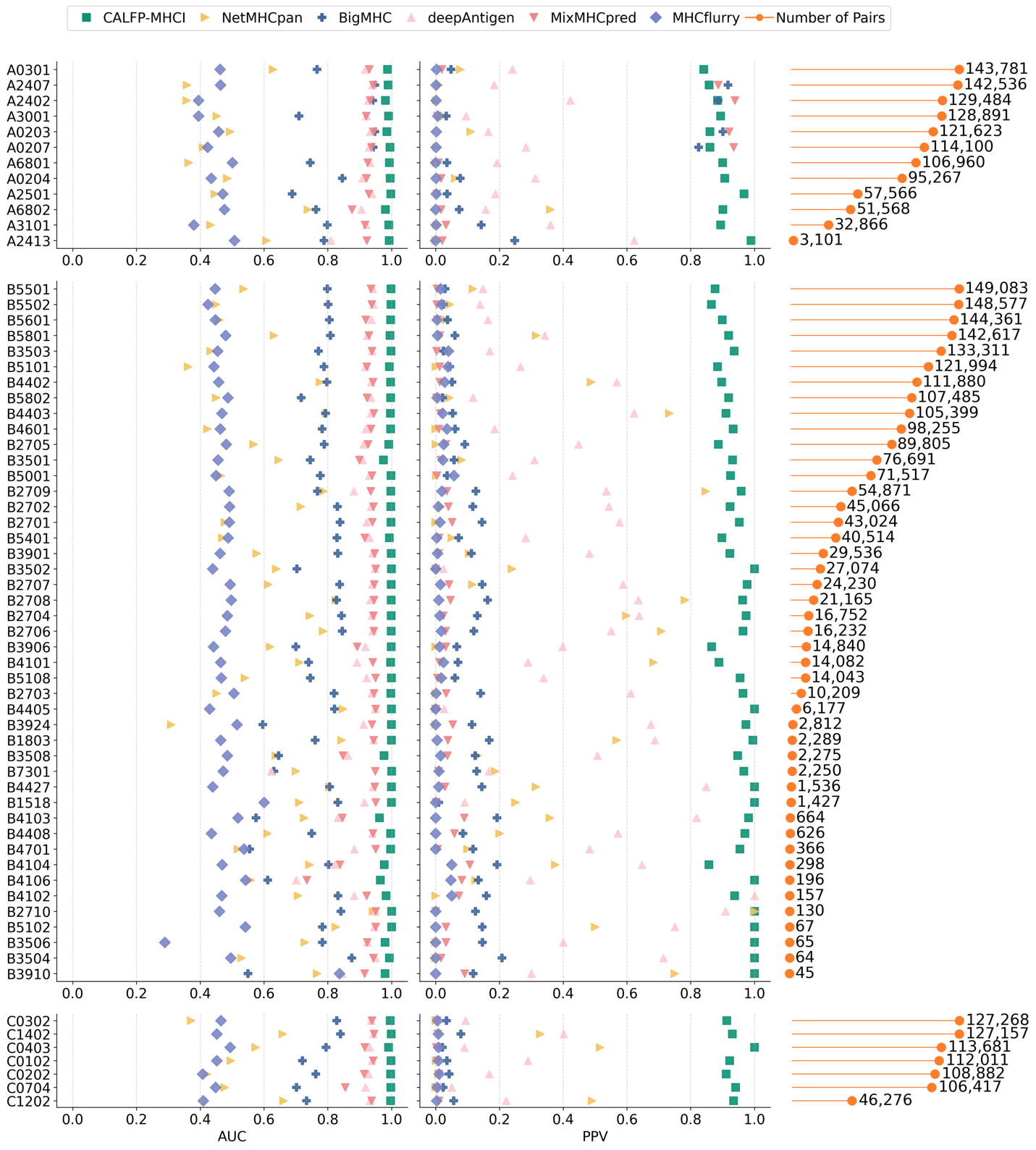
